# Epidermal activation of Hedgehog signaling establishes an immunosuppressive microenvironment in basal cell carcinoma by modulating skin immunity

**DOI:** 10.1101/768796

**Authors:** Sandra Grund-Gröschke, Daniela Ortner, Antal B. Szenes-Nagy, Nadja Zaborsky, Richard Weiss, Daniel Neureiter, Martin Wipplinger, Angela Risch, Peter Hammerl, Richard Greil, Maria Sibilia, Iris K. Gratz, Patrizia Stoitzner, Fritz Aberger

## Abstract

Genetic activation of Hedgehog (HH)/GLI signaling causes basal cell carcinoma (BCC), a very frequent non-melanoma skin cancer. Small molecule targeting of the essential HH effector Smoothened (SMO) proved an efficient medical therapy of BCC, although lack of durable responses and frequent development of drug resistance pose major challenges to anti-HH treatments. In light of the recent breakthroughs in cancer immunotherapy, we analyzed in detail the possible immunosuppressive mechanisms in HH/GLI-induced BCC. Using a genetic mouse model of BCC, we identified profound differences in the infiltration of BCC lesions with cells of the adaptive and innate immune system. Epidermal activation of HH/GLI led to an accumulation of immunosuppressive regulatory T cells, and to an increased expression of immune checkpoint molecules including PD-1/PD-L1. Anti-PD1 monotherapy, however, did not reduce tumor growth, presumably due to the lack of immunogenic mutations in common BCC mouse models, as shown by whole-exome sequencing. BCC lesions also displayed a marked infiltration with neutrophils, the depletion of which unexpectedly promoted BCC growth. The results provide a comprehensive survey of the immune status of murine BCC and provide a basis for the design of efficacious rational combination treatments. This study also underlines the need for predictive immunogenic mouse models of BCC to evaluate *in vivo* the efficacy of immunotherapeutic strategies.

## Introduction

Basal cell carcinoma (BCC) of the skin represents a very common human cancer entity with around 3-4 million new cases diagnosed per year in the US alone (Sekulic and Von Hoff, 2016). The etiology of BCC is based on the persistent ligand-independent activation of Hedgehog/Glioma-associated oncogene homolog (HH/GLI) signaling, caused in the vast majority of cases by inactivating mutations in the HH receptor and pathway repressor Patched-1 (PTCH1) or in fewer cases by activating mutations in the essential HH effector Smoothened (SMO). Irreversible activation of HH signaling culminates in high levels of oncogenic GLI transcription factors, which initiate and promote tumor growth by continuous hyperactivation of HH target genes involved in proliferation, survival, angiogenesis, stemness and (de-) differentiation (Epstein, 2008, Grachtchouk et al., 2000, Hahn et al., 1996, Hui and Angers, 2011, Johnson et al., 1996, Kasper et al., 2006, Nilsson et al., 2000, Oro et al., 1997, Ruiz i Altaba et al., 2007, Teglund and Toftgard, 2010, Xie et al., 1998). Although most BCC lesions are routinely removed by surgical excision, unresectable locally advanced and metastatic BCC require drug therapy. Small molecule inhibitors targeting SMO such as vismodegib and sonidegib have shown therapeutic efficacy in locally advanced and metastatic BCC, with overall response rates of 40-60 percent and complete responses in about 20 percent of patients (Atwood et al., 2014, Axelson et al., 2013, Casey et al., 2017, Dlugosz et al., 2012, Sekulic et al., 2017). However, despite the striking therapeutic efficacy of SMO inhibitors, their successful clinical use is limited and challenged by frequent *a priori* and acquired drug resistance, lack of durable responses, severe adverse effects and relapse of patients upon drug withdrawal. These limitations call for novel therapeutic regimens improving the response rates and durability of the therapeutic effect of HH inhibitors (Atwood et al., 2013, Atwood et al., 2015, Sekulic and Von Hoff, 2016, Sharpe et al., 2015, Tang et al., 2012, Whitson et al., 2018).

The recent breakthroughs in cancer immunotherapy that are based on our present understanding of how cancer cells evade the surveillance machinery of the adaptive and innate immune system (Chen and Mellman, 2013) have guided and paved the way to more efficacious, durable and even curative cancer therapies. For instance, therapeutic antibodies targeting immune checkpoints such as programmed death-1/programmed death-ligand 1 (PD1/PD-L1) signaling have been shown to re-instate the anti-tumoral immune response resulting in yet unprecedented therapeutic efficacy even in metastatic diseases (Okazaki et al., 2013, Pardoll and Allison, 2004, Wei et al., 2018). Intriguingly, two reports have already demonstrated a therapeutic benefit of anti-PD-1 treatment for BCC patients and a combination of vismodegib and pembrolizumab is currently evaluated in a clinical trial (trial ID: NCT02690948)(Chang et al., 2019, Lipson et al., 2017). Together, these data raise the hope that rational combination treatments targeting oncogenic HH/GLI and immunosuppressive mechanisms will synergistically improve the efficacy and durability of the therapeutic response of BCC patients with advanced or metastatic disease. A detailed understanding of the immune microenvironment of BCC as well as of the molecular player involved in establishing immune evasion is, therefore, of critical importance for the advancement of combination immunotherapy for unresectable BCC.

In this study, we strived to systematically investigate the molecular and cellular status of the immune microenvironment of HH/GLI-induced BCC, using a mouse model mimicking the genetic etiology of human BCC. We demonstrate that epidermal activation of HH/GLI signaling entails a variety of immunomodulatory mechanisms known to suppress immune recognition and subsequent eradication of neoplastic cells, thereby providing a basis for future combination treatments including immunotherapeutic drugs. In addition, we performed genomic sequencing of murine BCC-like lesions, which unlike in human BCC revealed no significant mutational burden, underlining the high need for predictive immunogenic murine BCC models to evaluate *in vivo* the efficacy of novel combinatorial immunotherapeutic treatments.

## Results

### Oncogenic HH/GLI signaling alters T cell populations in mouse BCC-like skin

The patients’ response to cancer immunotherapy correlates with high mutational burden of the cancer tissue, presumably due to increased immunogenicity as a consequence of neoantigen expression (Efremova et al., 2017, Samstein et al., 2019). Intriguingly, genomic sequencing of human BCC revealed that HH/GLI-driven skin cancers display an exceptionally high mutational burden with an average of 65 mutations per megabase (Bonilla et al., 2016), suggesting that BCC is likely to represent an immunogenic cancer entity. We, therefore, hypothesize that oncogenic HH/GLI signaling drives BCC growth also by suppressing the anti-tumoral immune response and possibly, also by recruiting inflammatory cells with tumor-promoting function. Although single case and prove-of-concept studies have suggested that immunotherapeutic approaches can be successful (Chang et al., 2019, Lipson et al., 2017), very little is known about how oncogenic HH/GLI signaling regulates tumor immunity in BCC. To investigate the immunological modulation of BCC-like tumors in mouse skin in response to ligand-independent Hh/Gli activation, we treated eight week old *K14CreER*^T^;*Ptch*^*f/f*^ (*Ptch*^*Δep*^) mice with tamoxifen (TAM) to induce irreversible Hh/Gli activation and BCC-like tumor formation, respectively. TAM-treated, Cre negative animals (*Ptch^f/f^*, controls) were used as controls (Fig. 1a). In agreement with previous reports (Biehs et al., 2018, Sternberg et al., 2018), mice with epidermal-specific deletion of Ptch1 displayed a hyperproliferative BCC-like phenotype that was most prominently observable on the ears (Fig. 1b).

**Figure 1:**
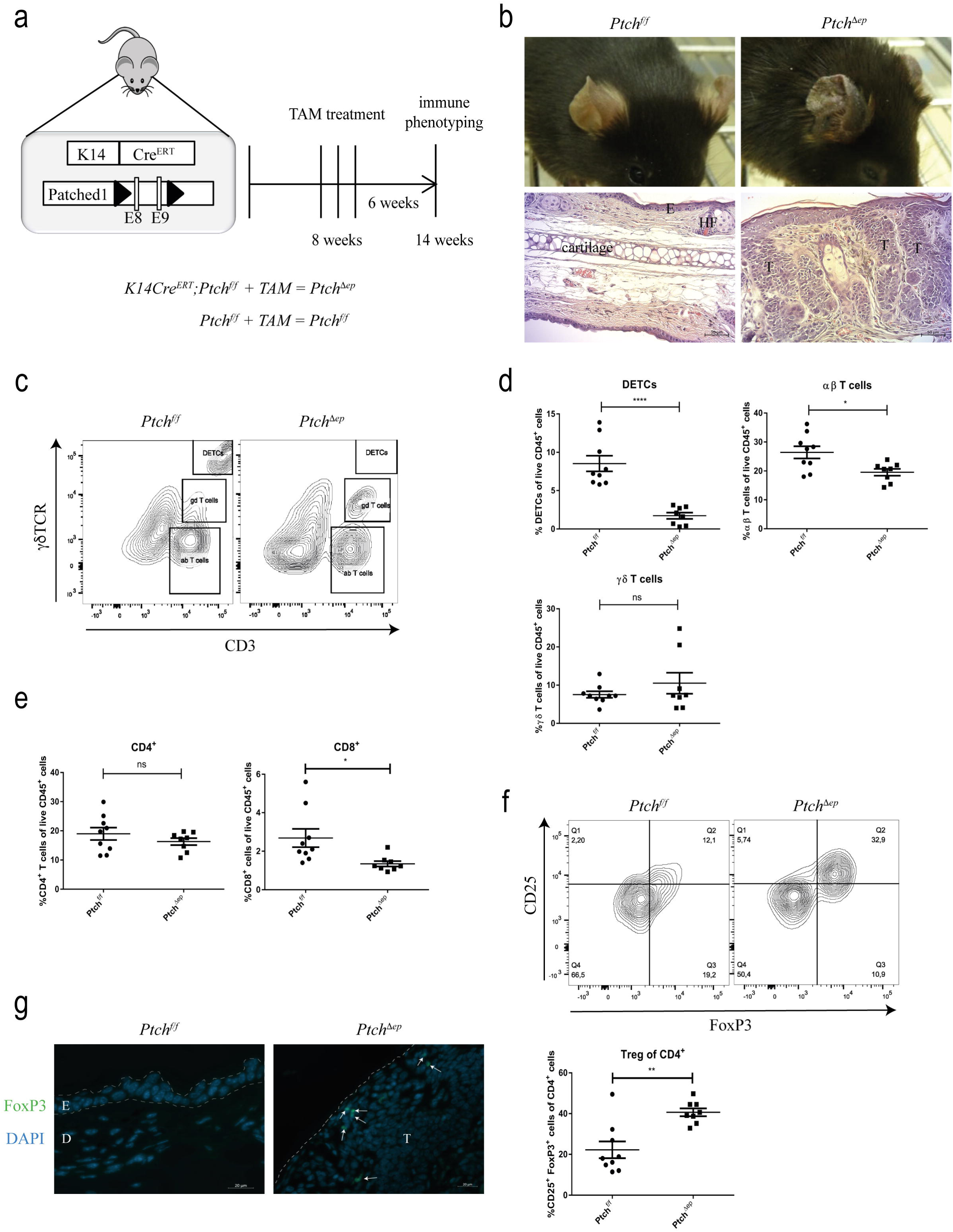
Oncogenic HH signaling leads to altered T cell populations in BCC-like skin. (a) Schematic illustration of the *K14CreER*^*T*^;*Ptch*^*f/f*^ (*Ptch*^*Δep*^) mouse model and TAM treatment schedule. (b) Representative figures of the phenotype (upper panel) and H&E stainings (lower panel) of *Ptch*^*f/f*^ and *Ptch*^*Δep*^ mouse ears. (c) Representative flow cytometry plots for T cell separation in mouse skin, using antibodies against γδTCR and CD3. (d, e and f) Flow cytometry analysis of (d) T cell populations, (e) CD4^+^ and CD8^+^ T cells and (f) CD25^+^ FoxP3^+^ Treg cells in *Ptch*^*Δep*^ and *Ptch*^*f/f*^ mouse skin. (f) Representative flow cytometry plot of CD25^+^ FoxP3^+^ Treg in mouse skin (upper panel) and quantitative result of the flow cytometry analysis (lower panel). (g) Immunofluorescence analysis of mouse ear for FoxP3 (green) and DAPI (blue). The arrows indicate the FoxP3^+^ cells. E: epidermis, D: dermis, T: tumor mass, HF: hair follicle.

To analyze alterations of adaptive immune cells in tumor lesions of *Ptch*^*Δep*^mice we performed flow cytometry analysis of skin T cell populations. Viable CD45^+^ cells were gated for CD3 versus γβTCR, resulting in three distinct T cell populations with CD3^+^ γδTCR^−^ cells representing αβ-T cells, CD3^+^ γδTCR^+^ cells representing γδ-T cells and cells that showed very high expression of both markers known as dendritic epidermal T cells (DETCs) (Fig. 1c). Interestingly, αβ-T cells and DETCs were decreased in *Ptch*^*Δep*^ mice in comparison to TAM-treated control mice, whereas the γδ-T cells remained unchanged (Fig. 1d). We validated these results by analyzing *K5cre^ER^*;*SmoM2 (SmoM2*) BCC mice (Eberl et al., 2012) expressing constitutively active oncogenic SMO in the epidermis after TAM injection. As shown in supplementary Figure S1a, CD3^+^ and CD3^high^ (DETC) T cells were also reduced in lesional skin of *SmoM2* mice compared to controls (suppl. Fig. S1a).

Having shown that αβ-T cells were decreased in skin tumors, we next addressed whether this applies to CD8^+^ or CD4^+^ T subsets. Gating for both markers revealed that cytotoxic CD8^+^ T cells were slightly lowered in the skin of tumor-bearing mice, whereas the CD4^+^ T cell population stayed unaffected (Fig. 1e). Of note, this CD8 decrease could be confirmed in *SmoM2* BCC mice. However, in contrast to *Ptch*^*Δep*^ mice CD4^+^ T cells were increased in *SmoM2* mice (suppl. Fig. S1b).

Regulatory T cells (Treg) cells play a pivotal role in cancer immune evasion, by suppressing the anti-tumoral immune response through different mechanisms such as the production of immunosuppressive cytokines (Mougiakakos et al., 2010, Takeuchi and Nishikawa, 2016). Although the percentages of CD4^+^ lymphocytes were not changed in *Ptch*^*Δep*^ mice, we asked whether FoxP3^+^ CD25^+^ Treg cells were altered. Notably, and as shown in Fig. 1f, FoxP3^+^ Tregs were strongly increased in mouse skin tumors compared to non-lesional control skin. To determine the localization of Treg cells, we performed immunofluorescence staining of *Ptch*^*Δep*^ mouse ears. We found Treg cells to be localized in intra- and peri-tumoral regions, suggesting a possible role of Tregs in immunosuppression of the BCC microenvironment (Fig. 1g).

### Increased immune checkpoint expression in mouse BCC-like skin

Having shown increased levels of immunosuppressive Tregs in mouse BCC-like skin, we next addressed the question of whether additional immunoregulatory mechanisms may support tumor immune escape. To this end, we performed a systematic screen for known immune checkpoints (Greil et al., 2017) via quantitative PCR of mouse BCC-like and control skin. Interestingly, we found *Pd-1* and its ligands *Pd-l1* and *Pd-l2* as well as *Tigit*, *Tim3* and *Cd226* to be significantly upregulated in skin tumors, whereas *Lag3* and *Cd96* were not significantly changed compared to control skin (Fig. 2a). In addition, we confirmed upregulation of *Pd-l1* mRNA levels in tumor lesions of *SmoM2* mice (suppl. Fig. S1c).

**Figure 2:**
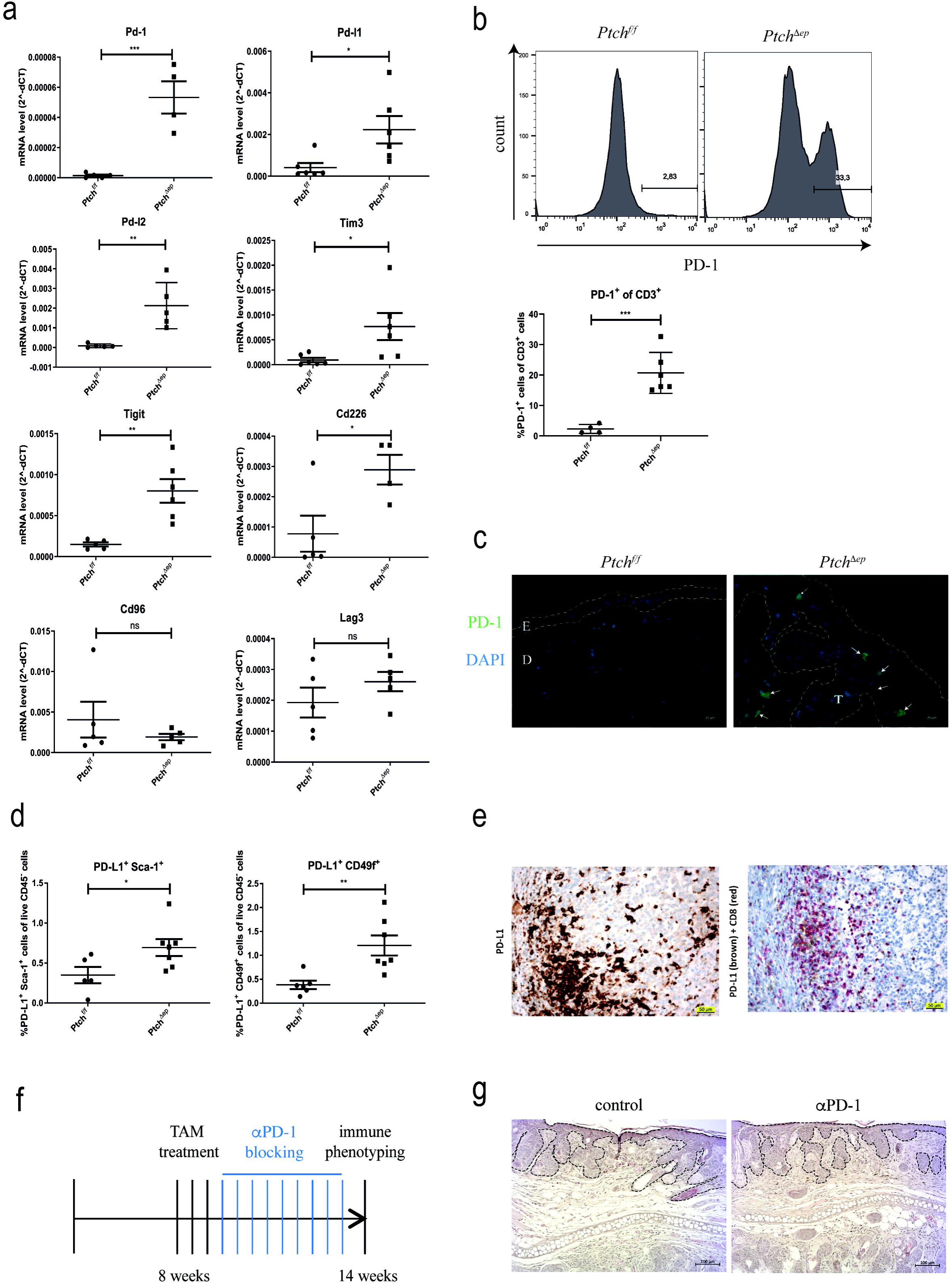
Increased immune checkpoint expression in BCC tumors. (a) qPCR analysis of mouse skin. (b) Representative flow cytometry plot showing PD-1 expression in mouse skin (upper panel) and flow cytometry analysis of PD-1 expression on CD3^+^ T cells (lower panel). (c) Immunofluorescence analysis of murine ear skin stained for PD-1 (green) and DAPI (blue). The arrows indicate PD-1^+^ cells. (d) Flow cytometry analysis of PD-L1 expression on CD45^−^ Sca-1^+^ and CD49f^+^ keratinocytes in mouse skin. (e) Immunohistochemical staining of human BCC skin sections for PD-L1 (brown) and CD8 (blue). (f) Treatment schedule of PD-1 blocking in previously TAM treated *K14CreER*^*T*^;*Ptch*^*f/f*^ BCC mice. (g) Representative images of H&E staining of ear skin of *Ptch*^*Δep*^ mice treated with anti-PD-1 blocking antibody or untreated (n=4 per group). The dashed lines mark the tumor area.

Given the clinical relevance of the PD-1/PD-L1 axis in cancer immunotherapy (Borghaei et al., 2015, Brahmer et al., 2015, Greil et al., 2017, Postow et al., 2015, Robert et al., 2015), we analyzed PD-1 expression on skin T cells of *Ptch*^*Δep*^ and control mice. As shown in Figure 2b, 20% of all T cells expressed PD-1 in tumor lesions of *Ptch*^*Δep*^ mice, whereas these cells were virtually absent in control skin (Fig. 2b). To determine the *in situ* localization of PD-1 positive cells in the skin, we performed immunofluorescence staining of mouse ears. We found most PD-1^+^ cells to be located adjacent or within the epithelial tumor compartment (Fig. 2c). Intriguingly, expression of the PD-1 ligand PD-L1 was mainly found in the Sca-1+/CD49f+ epithelial compartment of BCC-like lesions, suggesting an induction of this immune inhibitory molecule in response to Hh/Gli activation in keratinocytes (Fig. 2d), consistent with a recent study showing PD-L1 activation in a murine organoid model of Gli2-driven gastric cancer (Chakrabarti et al., 2018).

To show the relevance of these murine data to human pathophysiology, we performed immunohistochemical staining of human BCC specimen for CD8 and PD-L1. Human BCC present a pronounced infiltration with CD8^+^ T cells together with focal PD-L1 expression on tumor cells in close proximity to CD8^+^ T lymphocytes, suggesting inhibition of cytotoxic T cells by PD-L1 expressing BCC cells via immune checkpoint signaling. We also observed PD-L1 staining directly on CD8^+^ T lymphocytes (Fig. 2e).

Having shown that Pd-1 is upregulated on skin T cells of *Ptch*^*Δep*^ mice, we set out to test whether re-activation of the anti-tumor immune response via anti-PD-1 immune checkpoint inhibition results in tumor regression. For this purpose, we injected TAM-treated *K14CreER^T^;Ptch^f/f^* with anti-PD-1 blocking antibodies (Fig. 2f). However, and unlike in human case studies, anti-Pd-1 treatment was unable to reduce the tumor burden of *Ptch*^*Δep*^ mice (Fig. 2g). Analysis of the immune infiltrate of the skin via flow cytometry did not reveal significant alterations in the immune cells due to PD-1 therapy (suppl. Fig. S2). Since the response to immune checkpoint inhibition correlates with mutational burden (Samstein et al., 2019), we hypothesized that the lack of sufficient immunogenic mutations in our murine model of BCC may account for the failure of PD-1 therapy. Therefore, we performed whole exome sequencing (complete mouse exome coverage 49.6 Mb) of mouse BCC-like lesions (n=3) and control skin (n=3) to survey the mutational landscape of *Ptch*^*Δep*^ tumor lesions. As shown in Table S1, we were unable to identify any mutations in two of three mice. One mouse tumor showed two mutations with unknown immunological consequence (suppl. Table S1). Furthermore, analysis of the copy number variations revealed minor negligible chromosomal alterations in all three mice analyzed (suppl. Fig. S3). Based on these data, we propose that *Ptch*^*Δep*^ tumors do not express neoantigens and thus, are non-immunogenic. This underlines the need for next-generation immunogenic mouse models of HH/GLI-driven skin cancer to evaluate immunotherapeutic strategies.

### Mouse BCC-like skin displays an altered cytokine and chemokine expression profile

Having shown that largely unmutated *Ptch*^*Δep*^ lesions comprise altered T cell populations in comparison to *Ptch*^*f/f*^ skin, we hypothesized that also the cytokine and chemokine expression profiles differ between tumor lesions and control mice. We, therefore, performed by qPCR analysis and Luminex profiling for systematic mRNA and protein expression analysis of tumor and control skin for known immune-modulating cytokines and chemokines (for summary see Table 1). Of note, we found increased levels of several immunosuppressive cytokines such as *Tslp*, *Tgfβ*,*Il10* and *Inos* as well as the pro-inflammatory cytokines as *Tnfα*, *Il1β*, *Ifnγ* and *Il17* in tumor lesions compared to control skin. Furthermore, BCC-like lesions expressed elevated levels of immune-cell attracting chemokines such as *Ccl2*, *Ccl3* and *Ena78* (Gaga et al., 2008, Griffith et al., 2014). We observed similar changes in cytokine/chemokine expression profiles in *SmoM2* BCC mice (suppl. Fig. S4). We conclude that persistent Hh/Gli signaling in epidermal cells induces pronounced changes in the expression of immune-modulatory and chemoattractant factors that contribute to the formation of an immunosuppressive tumor microenvironment, including shifts of T lymphocyte and possibly also myeloid cell populations.

**Table 1:** Cytokine and chemokine expression profiles in murine BCC

| **Table 1: Cytokine and chemokine expression profiles in murine BCC** |  |  |  |  |
| --- | --- | --- | --- | --- |
| **Cytokines** | **mRNA** |  | **Protein** |  |
| **Gene name** | **fold change** | **p-value** | **fold change** | **p-value** |
| Il10 | 4.2 | 0.0523 | nd | nd |
| Tgfβ | 4.5 | 0.0039 | nd | nd |
| Il1β | 13.4 | 0.0024 | 3.4 | 0.1036 |
| Ifnγ | 7.8 | 0.0011 | nd | nd |
| Il17 | 27.5 | 0.0098 | nd | nd |
| Gmcsf | 3.9 | 0.0104 | 1 | 0.9949 |
| Inos | 5.6 | 0.0425 | nd | nd |
| Cox2 | 2.2 | 0.0747 | nd | nd |
| Tslp | 47.3 | 0.0710 | 2.6 | 0.0010 |
| Tnf | 2.2 | 0.0688 | 2.6 | 0.0083 |
| **Chemokines** | **mRNA** |  | **Protein** |  |
| **Gene name** | **fold change** | **p-value** | **fold change** | **p-value** |
| Mip-2 | nd | nd | 1.2 | 0.6193 |
| Ena78 | 13.1 | 0.0003 | 1.4 | 0.4529 |
| Gro-α | nd | nd | 1.5 | 0.3141 |
| Ccl2/Mcp-1 | 17 | 0.0418 | 4.6 | 0.0284 |
| Ccl3/Mip-1α | 3.8 | 0.0125 | 4.1 | 0.0161 |
| Ccl7/Mcp-3 | nd | nd | 2.5 | 0.2955 |
| Cxcl10/Ip-10 | nd | nd | 1.1 | 0.9148 |

nd = not determined

### Oncogenic HH signaling leads to altered innate immunity in mouse BCC-like skin

Having demonstrated that Hh/Gli signaling in mouse BCC-like tumors results in a significant increase in chemokine levels such as Ccl2 and Ccl3 – two well-known chemoattractants for innate immune cells (Deshmane et al., 2009, Griffith et al., 2014) - we next analyzed quantitatively and qualitatively by flow-cytometry analysis, whether epidermal Hh pathway activation results in changes of the innate immune cell population. To this end, we gated myeloid cells in skin samples of *Ptch*^*f/f*^ and *Ptch*^*Δep*^ mice by their CD11b and CD11c expression. Intriguingly, we found a pronounced increase in CD11b^+^ cells in mouse skin tumors (Fig. 3a), which we confirmed also in *SmoM2* mice. However, *SmoM2* mice showed a strong decrease in CD11c expression (suppl. Fig. S5a), while we did not see this prominent effect in *Ptch*^*Δep*^ mice (Fig. 3a).

**Figure 3:**
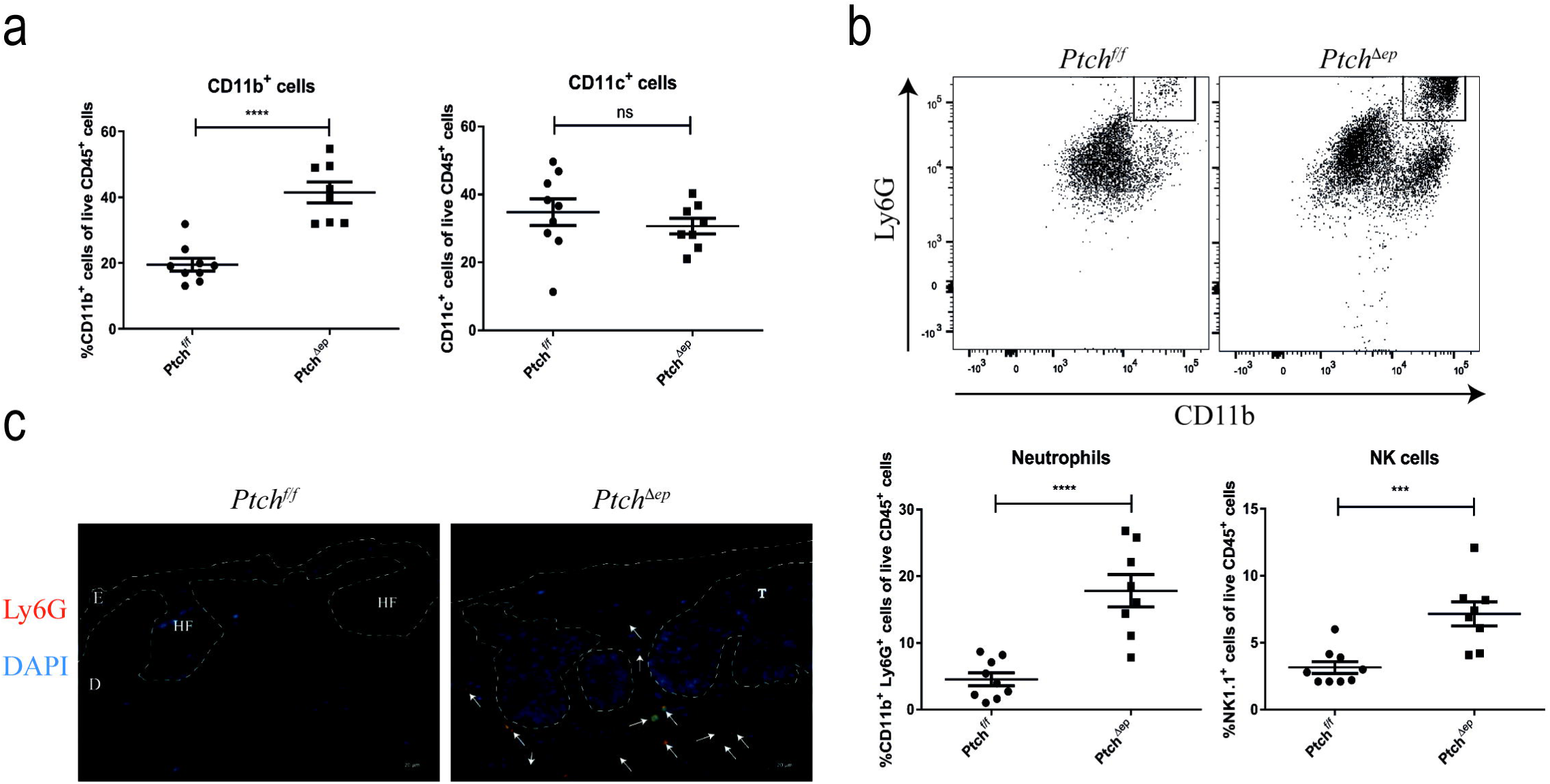
Oncogenic HH signaling leads to altered innate immunity in BCC. (a) Flow cytometry analysis of murine skin for CD11b and CD11c expression in *Ptch*^*Δep*^ and *Ptch*^*f/f*^control mice. (b) Representative flow cytometry analysis plot for expression of Ly6G and CD11b (upper panel) and flow cytometry analysis for Ly6G and NK1.1 expression (lower panel). (c) Immunofluorescence staining of murine ear skin for Ly6G (red) and DAPI (blue). The arrows indicate Ly6G^+^ cells. E: epidermis, D: dermis, T: tumor mass, HF: hair follicle.

Further analysis of CD11b^+^ cells using the neutrophil marker Ly6G and the NK cell marker NK1.1 revealed a noticeable recruitment of both innate immune cell populations in *Ptch*^*Δep*^ skin lesions (Fig. 3b). Ly6G^+^ cells were also increased in skin BCC-like lesions of *SmoM2* mice (suppl. Fig. S5b). To better understand the putative role of neutrophils in BCC-like lesions, we first performed immunofluorescence staining of Ly6G^+^ cells in skin samples of *Ptch*^*f/f*^ and *Ptch*^*Δep*^ mice. We found most of the Ly6G^+^ cells to be located in the peri-tumoral region with only rare cells within the tumor nest itself (Fig. 3c).

Since Fan et al. have provided evidence that in *SmoM2* BCC mice Ly6G^+^ CD11b^+^ cells represent MDSCs with immunosuppressive function (Fan et al., 2014), we analyzed Ly6G^+^/CD11b^+^ cells of *Ptch*^*Δep*^ mice for the MDSC marker arginase (for antibody validation see suppl. Fig. S6). As shown in Fig. 4a, Ly6G^+^/CD11b^+^ cells of *Ptch*^*Δep*^ mice did not express the MDSC marker arginase, suggesting that tumor lesions of *Ptch*^*Δep*^ mice are infiltrated by neutrophils rather than MDSCs. To shed light on the role of tumor-infiltrating neutrophils in BCC, we next depleted *Ptch*^*Δep*^ mice of neutrophils by injecting anti-Ly6G depletion antibody (clone 1A8)(Fig. 4b). We proved successful depletion of neutrophils in the skin by flow cytometry (Fig. 4c). Intriguingly, depletion of neutrophils enhanced the growth of tumor lesions in *Ptch*^*Δep*^ mice (Fig. 4d and 4e), suggesting that neutrophils, known for their tumor-promoting function in a variety of malignant entities (Galdiero et al., 2018), may constitute an as yet unknown component of the anti-tumoral immune response in BCC. Future studies are required to better understand the possible anti-cancer activity of neutrophils in the context of HH/GLI-driven BCC.

**Figure 4:**
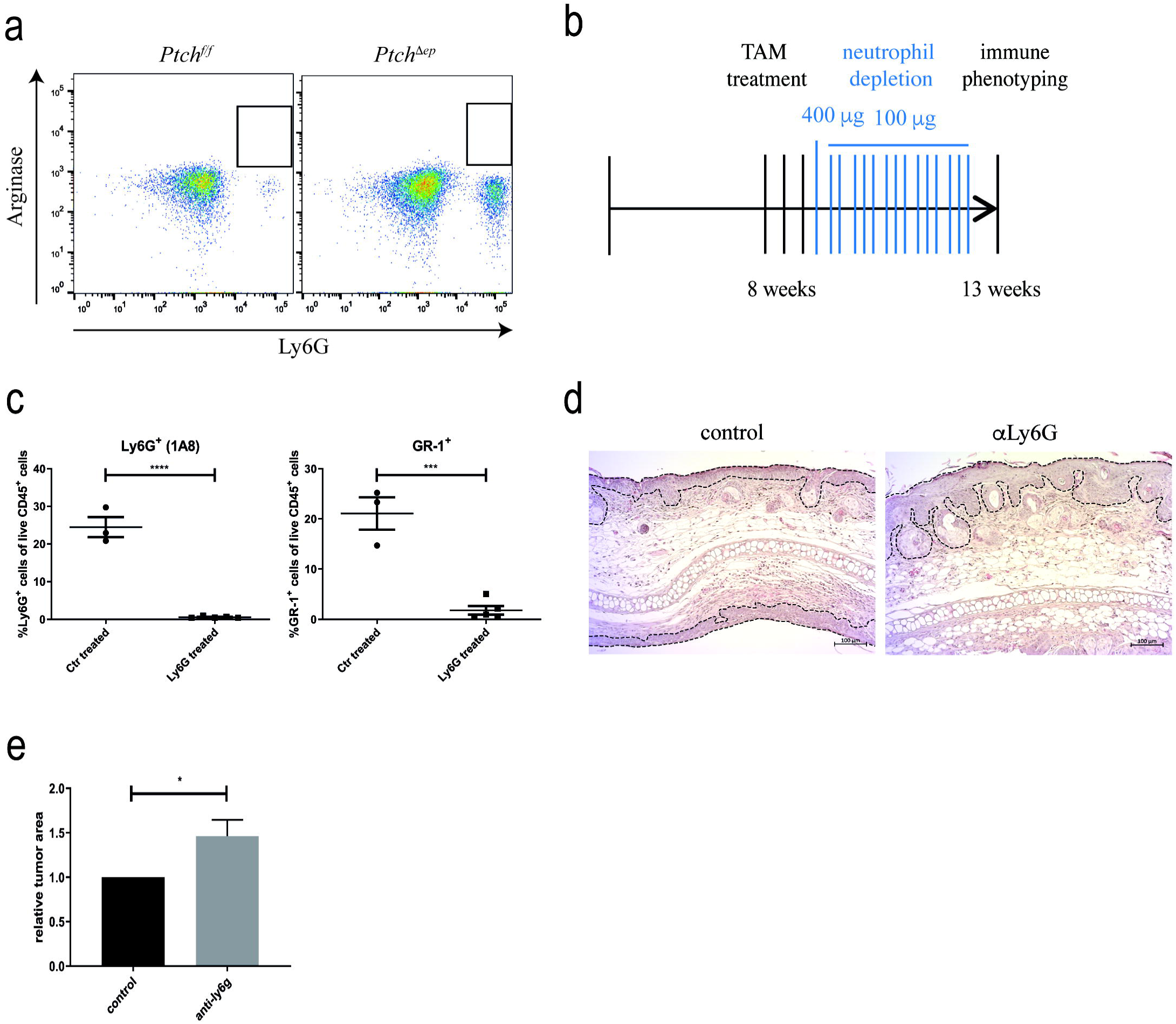
Neutrophil depletion in BCC mice accelerates tumor growth. (a) Representative flow cytometry analysis plots of arginase staining in *Ptch*^*Δep*^ and *Ptch*^*f/f*^ mouse skin. (b) Treatment schedule for neutrophil depletion in *K14CreER*^*T*^;*Ptch*^*f/f*^ mice previously injected with TAM. (c) Flow cytometry analysis of murine skin for Ly6G or Gr-1 expression verifying efficient depletion of neutrophils. (d) Representative H&E staining comparing tumor development in ear skin of *Ptch*^*Δep*^ mice not depleted (left) or depleted (right) of neutrophils during BCC progression. The dashed lines mark the tumor area. (e) Quantification of the tumor area in the ear skin of *Ptch*^*Δep*^ mice not depleted (control, n=3) or depleted (anti-Ly6G, n=5) of neutrophils during BCC progression.

## Discussion

The continuous growth and spread of malignant cells depend on the generation of an immunosuppressive microenvironment to prevent an efficient anti-tumoral immune response. The restoration of the immune response, for instance by the use of checkpoint inhibitors has marked a breakthrough in cancer therapy with yet unprecedented success in responding patients (Wei et al., 2018). A high mutational burden of cancer cells has been shown to correlate with the clinical response and success of immunotherapy using checkpoint inhibitors (Samstein et al., 2019). Given the exceptionally high mutational burden of human BCC (Bonilla et al., 2016), we here hypothesized that i) HH/GLI-induced BCC are immunogenic, ii) HH/GLI signaling establishes an immune-suppressive immune microenvironment and iii) HH/GLI-induced BCC are susceptible to immunotherapy. To investigate in detail the poorly defined role of HH/GLI in the process of immune evasion, we used genetic mouse models of BCC that closely mimic the genetics of driver mutations in human BCC. In the present study, we show that active epidermal HH/GLI signaling is a potent inducer of immunosuppressive mechanisms such as the accumulation of Tregs, the upregulation of immune checkpoints and the production of immunosuppressive growth factors and cytokines including IL10 and TGFβ within the tumor lesions. Although we did not determine the *in vivo* source of these immunosuppressive signals, it is well possible that these key immunosuppressive factors derive directly from cancer cells in response to active HH/GLI. This is supported by previous studies showing that SHH/GLI signaling can induce IL10 expression in a murine model of colitis and pancreatitis. Similarly, overexpression of active GLI2 in T cells can cell-autonomously enhance IL10 and TGFβ expression (Furler and Uittenbogaart, 2012, Lee et al., 2016, Papaioannou et al., 2019, Zhou et al., 2012). In line with the expression of IL10 and TGFβ, we show increased numbers of Treg cells in *Ptch*^*Δep*^ tumors, similar to human BCC, where Omland et al. provided evidence that active HH/GLI signaling can induce Treg accumulation along with a strong increase of TGFβ expression (Omland et al., 2016). Furthermore, Hanna et al. have recently shown that HH/GLI signaling can drive Treg recruitment in breast cancer, while pathway inhibition reduced the number of Tregs in the tumor (Hanna et al., 2019). The high prevalence of Tregs within the tumor microenvironment is, thus, likely to constitute a major mechanism of immunosuppression by HH/GLI signaling in cancer.

In addition to an increased level of Treg cells, we demonstrated enhanced expression of the immune checkpoint molecules PD-1 and its cognate ligands PD-L1 and PD-L2 in *Ptch*^*Δep*^ mouse tumors, with PD-L1 expressed preferentially on epithelial tumor cells and PD-1 on adjacent CD8^+^ T cells. This suggests that HH/GLI signaling in BCC can induce PD-L1 expression on tumor cells and thereby, contribute to the suppression of infiltrating cytotoxic T cells directed against the tumor cells. Together with previous reports on the role HH/GLI in gastric cancer showing PD-L1 upregulation by HH/GLI signaling (Chakrabarti et al., 2018, Holla et al., 2016), our findings in BCC suggest, that HH/GLI-mediated induction of the PD-L1/PD-1 immune checkpoint represents a more general mechanism of HH-mediated immune escape in cancer. Intriguingly, the increased expression in *Ptch*^*Δep*^ lesions of additional immune checkpoints such as Tigit and Tim3 (Fig. 2a)(Greil et al., 2017) point to a broader involvement of oncogenic HH/GLI signaling in the suppression of cytotoxic T cell responses, increasing the therapeutic opportunities for combination treatments with anti-HH and immune checkpoint inhibitors.

In agreement with our findings on immune checkpoint expression profiles of murine BCC-like lesions, first proof-of-concept studies support a therapeutic benefit of anti-PD-1 treatment for patients with advanced or metastatic and SMO inhibitor resistant BCC (Chang et al., 2019, Ikeda et al., 2016, Lipson et al., 2017). These positive initial results underline the importance of predictive mouse models of BCC that allow the evaluation of innovative combination therapies including immunotherapeutics. Thus, our characterization of the immunological alterations in response to Hh/Gli activation in *Ptch*^*Δep*^ and *SmoM2* mouse models is crucial to understand the immune infiltration and cytokine/chemokine expression profiles in BCC-like skin lesions. It proves the usefulness of these mouse models to study in detail the pathologically relevant changes in the immune microenvironment of HH/GLI induced BCC. However, the failure of *Ptch*^*Δep*^ mice to respond to anti-PD-1 immune checkpoint inhibitor therapy, which may be due to the absence of immunogenic mutations as shown by whole-exome sequencing, reveals the limitations of this common BCC mouse model for immunotherapy approaches. The development of immunogenic murine BCC models is, therefore, paramount for future preclinical studies.

In addition to changes of the T cell populations, we also identified pronounced infiltration of BCC-like lesions by Ly6G^pos^/Arginase^neg^ neutrophils, which was in agreement with the elevated expression of chemokines such as Ccl2 and Ccl3, two potent chemoattractants for myeloid cells (Deshmane et al., 2009, Griffith et al., 2014). Since neutrophil depletion accelerated tumor development in *Ptch*^*Δep*^ mice, we conclude that in HH/GLI-driven skin cancer neutrophils, which we found to be mainly located in the tumor periphery, fulfill anti-tumorigenic functions. Although neutrophils are well known for their tumor-promoting function (Coffelt et al., 2015, Fridlender et al., 2009, Mantovani, 2009), evidence also suggests a critical contribution of neutrophils to the anti-tumoral immune response, reflecting their remarkable plasticity. For instance, neutrophils have been shown to enhance MHC-I expression, thereby increasing the anti-tumoral response of adaptive immunity. Further, tumor-associated neutrophils can promote the proliferation of effector T cells and impede tumor progression by the release of pro-apoptotic TRAIL (reviewed in (Galdiero et al., 2018). Whether these mechanisms account for the tumor-suppressive activity of neutrophils in murine BCC skin needs to be addressed in future studies. In this context it is also noteworthy that Fan et al. have provided evidence that pre-natal epidermal activation of *SmoM2* resulted in BCC-like lesions enriched for Ly6G^pos^ pro-tumorigenic MDSCs. Recruitment of MDSCs was inhibited by systemic chemical inhibition of Ccl2 receptor, resulting in reduced tumor growth (Fan et al., 2014). However, in contrast to our data and studies on human BCC, *SmoM2* mice with pre-natal Hh/Gli activation and MDSC enrichment did not show enhanced Treg numbers, which may point to subtle differences in the immune modulation of the distinct mouse models used.

Taken together, we conclude that the mere genetic activation of HH/GLI signaling in epidermal cells induces profound changes within the immune microenvironment of BCC-like lesions, thereby establishing a potent immunosuppressive milieu, which is likely to inhibit the anti-tumoral immune response against BCC cells with high mutational burden. Therapeutic strategies directed against immunosuppressive mechanisms activated by HH/GLI including several checkpoint inhibitors warrant further evaluation, though for pre-clinical *in vivo* studies it has to be considered that in general the genetic mouse models commonly used in HH-related skin cancer research do not properly reflect the UV-induced high mutational burden and potential immunogenicity of human BCC. Our results also call for next generation, immunogenic models of BCC to explore the full therapeutic potential of treatments including HH pathway inhibitors in combination with selected immunotherapeutic drugs.

## Material and Methods

### Mice

*K14CreER*^*T*^ (#005107) and *Ptch*^*f/f*^ (#012457) mice were obtained from the Jackson Laboratory (Bar Harbor, ME) and genotyped according to the supplier’s instructions. For tumor induction 8-week old mice and Cre negative *Ptch*^*f/f*^ animals were orally administered three times with 1 mg TAM (Sigma, St Louis, MO) dissolved in 10% ethanol/corn oil. Mice were analyzed six weeks after treatment, when they showed prominent pigmented lesions on the ears (also see Fig. 1a and 1b). Detailed information on *SmoM2* mice is provided in the supplementary material. All studies were performed on mice of both genders.

For PD-1 blocking, TAM-treated *K14CreER*^*T*^;*Ptch*^*f/f*^ mice were intraperitoneally injected every third day for 5 weeks with 250 µg of anti-PD-1 blocking antibodies (clone RMP1-14, BioXCell, West Lebanon, NH), according to previously published papers (Curran et al., 2010, John et al., 2013, Mittal et al., 2014, Ngiow et al., 2015). A schematic overview is provided in Fig. 3a.

Neutrophils were depleted by intraperitoneal injection of anti-Ly6G antibodies (clone 1A8, BioXCell, West Lebanon, NH). After TAM treatment, *K14CreER*^*T*^;*Ptch*^*f/f*^ mice received an initial dose of 400 µg anti-Ly6G followed by thrice weekly injections of 100 µg for 4 weeks, according to Coffelt et al. (Coffelt et al., 2015). Successful depletion was verified weekly via flow cytometry staining of mouse blood. All studies were performed on mice of both genders.

### Flow cytometry

For flow cytometry analysis of *K14CreER*^*T*^;*Ptch*^*f/f*^ and control mice, tumor and control skin was digested for 45 min at 37°C with 2 mg/ml collagenase XI, 0.5 mg/ml hyaluronidase und 0.1 mg/ml DNase in DMEM (all from Sigma, St Louis, MO). Skin cell suspensions were filtered through 70 µm cell strainer (Corning, Corning, NY). Cells were stained with directly conjugated antibodies for 30 min at 4°C in the dark. Separation of dead cells was accomplished with fixable viability dye eFluor 780, eFluor 520 or eFluor 506 (dilution 1:1000, Thermo Fisher Scientific, Waltham, MA). To separate immune cells from other skin cells CD45 as pan-leukocyte marker was included in all skin panels. Cells were fixed by using the Foxp3/transcription factor staining buffer set (Thermo Fisher Scientific, Waltham, MA) followed by intracellular staining. All experiments were performed on the BD FACS Canto II (BD Biosciences, Franklin Lakes, NJ). For data analysis the BD FACSDiva™ and the FlowJo® software (Tree Star) were used. Antibodies used for flow cytometry are listed in suppl. Table S2.

### Quantitative PCR

Total RNA isolation and qPCR analysis of mRNA expression was carried out as describes previously (Eberl et al., 2012). The quantity and quality of isolated RNAs were assessed by Agilent Bioanalyzer 2200 system (Agilent Technologies, Santa Clara, CA). qPCR was done on Rotor-Gene Q (Qiagen, Hilden, Germany) by using the GoTaq (Promega, Madsion, WI) or the Luna (NEB, Ipswich, MA) qPCR master mix. Details about primers used in the study are listed in suppl. Table S3.

### Luminex cytokine profiling

Mouse dorsal skin was homogenized with the Ultra-Turrax® (IKA, Staufen, Germany) in PBS supplemented with protease inhibitor (Sigma, St Louis, MO) and filtered through Corning® Costar® Spin-X® Plastic Centrifuge Tubes (Sigma, St Louis, MO). For cytokine profiling the ProcartaPlex™ multiplex system (Thermo Fisher Scientific, Waltham, MA) was used. The array analyses were carried out according to the supplier’s instructions. Measurement was performed on the Magpix instrument (Luminex corp, Austin, TX) and the analysis was done with the ProcartaPlex Analyst 1.0 software (Thermo Fisher Scientific, Waltham, MA).

### Histological analysis and immunofluorescence

Mouse ears were fixed overnight in 4% paraformaldehyde before paraffin embedding. 4 µm-sections from paraffin-embedded tissue were prepared for hematoxylin and eosin and immunofluorescence staining. Before staining, samples were deparaffinized and sections were blocked for 1 hour in 1% BSA. For immunofluorescence staining, slides were incubated overnight with antibodies against CD8, FoxP3, PD-1 and Ly6G. For intracellular staining of FoxP3, Triton X-100 (Sigma, St Louis, MO) was added to a final concentration of 0.1%. Alexa 488- or Alexa 555-conjugated secondary antibodies were used for detection. Slides were mounted with Fluoroshield Mounting Medium with DAPI (Abcam, Cambridge, United Kingdom). Antibodies used for staining and detection are listed in suppl. Table S4. All pictures were taken on a Zeiss Axio Observer Z1 microscope (Carl Zeiss, Oberkochen, Deutschland) using ZEN 2.6 software (Carl Zeiss, Oberkochen, Deutschland).

For quantification of the tumor area after neutrophil depletion, representative pictures from ears of three wildtype and three tumor mice were quantified with the ImageJ software by three independent researchers in a blinded manner.

### Statistical analysis

Significant differences between two groups were determined using an unpaired two-tailed student-*t-*test. *p* values of < 0.05 were considered significant and *p* values have been labelled and designated as follows: * *p* < 0.05, ** *p* < 0.01, *** *p* < 0.001 and **** *p* < 0.0001. All values are given as means (+/-SEM) and were analyzed by GraphPad Prism® 8 (GraphPad, San Diego, CA).

### Ethics

Human BCC tissue arrays for immunohistochemistry analyses were used in accordance with the guidelines of the Austrian ethics committee. Animal studies were approved by the institution authorities and by the Federal Ministry of Science, Research and Economy (BMWFW-66.012/0016-WF/V/3b/2015 and BMWFW-66.011/0030-II/3b/2014).

## Acknowledgments

We dedicate this work to our dear colleague Peter Hammerl, who sadly passed away during the course of this study. This work was supported by the Austrian Science Fund (FWF project W1213), the priority program Allergy-Cancer-Bionano Research Center of the University of Salzburg, and the Cancer Cluster Salzburg research grant of the County of Salzburg. The authors are grateful to the animal caretakers, to Drs. Angelika Stöcklinger for continuous support, Walter Stoiber and Angela Zissler for methodical input, to Gernot Posselt and Silja Wessler for help with microscopy and to Mag. Suzana Tesanovic and Mag. Dominik Elmer for help with the preparation of animals. Furthermore, the authors would like to express special thanks to Georg Stockmaier for critical reading of the manuscript.

## References

Atwood SX, Li M, Lee A, Tang JY, Oro AE. GLI activation by atypical protein kinase C iota/lambda regulates the growth of basal cell carcinomas. Nature 2013;494(7438):484–8.

Atwood SX, Sarin KY, Whitson RJ, Li JR, Kim G, Rezaee M, et al. Smoothened variants explain the majority of drug resistance in basal cell carcinoma. Cancer Cell 2015;27(3):342–53.

Atwood SX, Whitson RJ, Oro AE. Advanced treatment for basal cell carcinomas. Cold Spring Harbor perspectives in medicine 2014;4(7):a013581.

Axelson M, Liu K, Jiang X, He K, Wang J, Zhao H, et al. U.S. Food and Drug Administration approval: vismodegib for recurrent, locally advanced, or metastatic basal cell carcinoma. Clin Cancer Res 2013;19(9):2289–93.

Biehs B, Dijkgraaf GJP, Piskol R, Alicke B, Boumahdi S, Peale F, et al. A cell identity switch allows residual BCC to survive Hedgehog pathway inhibition. Nature 2018;562(7727):429–33.

Bonilla X, Parmentier L, King B, Bezrukov F, Kaya G, Zoete V, et al. Genomic analysis identifies new drivers and progression pathways in skin basal cell carcinoma. Nat Genet 2016;48(4):398–406.

Borghaei H, Paz-Ares L, Horn L, Spigel DR, Steins M, Ready NE, et al. Nivolumab versus Docetaxel in Advanced Nonsquamous Non-Small-Cell Lung Cancer. The New England journal of medicine 2015;373(17):1627–39.

Brahmer J, Reckamp KL, Baas P, Crino L, Eberhardt WE, Poddubskaya E, et al. Nivolumab versus Docetaxel in Advanced Squamous-Cell Non-Small-Cell Lung Cancer. The New England journal of medicine 2015;373(2):123–35.

Casey D, Demko S, Shord S, Zhao H, Chen H, He K, et al. FDA Approval Summary: Sonidegib for Locally Advanced Basal Cell Carcinoma. Clin Cancer Res 2017;23(10):2377–81.

Chakrabarti J, Holokai L, Syu L, Steele NG, Chang J, Wang J, et al. Hedgehog signaling induces PD-L1 expression and tumor cell proliferation in gastric cancer. Oncotarget 2018;9(100):37439–57.

Chang ALS, Tran DC, Cannon JGD, Li S, Jeng M, Patel R, et al. Pembrolizumab for advanced basal cell carcinoma: An investigator-initiated, proof-of-concept study. J Am Acad Dermatol 2019;80(2):564–6.

Chen DS, Mellman I. Oncology meets immunology: the cancer-immunity cycle. Immunity 2013;39(1):1–10.

Coffelt SB, Kersten K, Doornebal CW, Weiden J, Vrijland K, Hau CS, et al. IL-17-producing gammadelta T cells and neutrophils conspire to promote breast cancer metastasis. Nature 2015;522(7556):345–8.

Curran MA, Montalvo W, Yagita H, Allison JP. PD-1 and CTLA-4 combination blockade expands infiltrating T cells and reduces regulatory T and myeloid cells within B16 melanoma tumors. Proc Natl Acad Sci U S A. 107; 2010. p. 4275–80.

Deshmane SL, Kremlev S, Amini S, Sawaya BE. Monocyte chemoattractant protein-1 (MCP-1): an overview. J Interferon Cytokine Res 2009;29(6):313–26.

Dlugosz A, Agrawal S, Kirkpatrick P. Vismodegib. Nature reviews Drug discovery 2012;11(6):437–8.

Eberl M, Klingler S, Mangelberger D, Loipetzberger A, Damhofer H, Zoidl K, et al. Hedgehog-EGFR cooperation response genes determine the oncogenic phenotype of basal cell carcinoma and tumour-initiating pancreatic cancer cells. EMBO molecular medicine 2012;4(3):218–33.

Efremova M, Finotello F, Rieder D, Trajanoski Z. Neoantigens Generated by Individual Mutations and Their Role in Cancer Immunity and Immunotherapy. Front Immunol 2017;8.

Epstein EH. Basal cell carcinomas: attack of the hedgehog. Nature reviews Cancer 2008;8(10):743–54.

Fan Q, Gu D, Liu H, Yang L, Zhang X, Yoder MC, et al. Defective TGF-beta signaling in bone marrow-derived cells prevents hedgehog-induced skin tumors. Cancer research 2014;74(2):471–83.

Fridlender ZG, Sun J, Kim S, Kapoor V, Cheng G, Ling L, et al. Polarization of tumor-associated neutrophil phenotype by TGF-beta: “N1” versus “N2” TAN. Cancer Cell 2009;16(3):183–94.

Furler RL, Uittenbogaart CH. GLI2 regulates TGF-beta1 in human CD4+ T cells: implications in cancer and HIV pathogenesis. PLoS One 2012;7(7):e40874.

Gaga M, Ong YE, Benyahia F, Aizen M, Barkans J, Kay AB. Skin reactivity and local cell recruitment in human atopic and nonatopic subjects by CCL2/MCP-1 and CCL3/MIP-1alpha. Allergy 2008;63(6):703–11.

Galdiero MR, Varricchi G, Loffredo S, Mantovani A, Marone G. Roles of neutrophils in cancer growth and progression. J Leukoc Biol 2018;103(3):457–64.

Grachtchouk M, Mo R, Yu S, Zhang X, Sasaki H, Hui CC, et al. Basal cell carcinomas in mice overexpressing Gli2 in skin. Nat Genet 2000;24(3):216–7.

Greil R, Hutterer E, Hartmann TN, Pleyer L. Reactivation of dormant anti-tumor immunity - a clinical perspective of therapeutic immune checkpoint modulation. Cell Commun Signal 2017;15(1):5.

Griffith JW, Sokol CL, Luster AD. Chemokines and chemokine receptors: positioning cells for host defense and immunity. Annu Rev Immunol 2014;32:659–702.

Hahn H, Wicking C, Zaphiropoulous PG, Gailani MR, Shanley S, Chidambaram A, et al. Mutations of the human homolog of Drosophila patched in the nevoid basal cell carcinoma syndrome. Cell 1996;85(6):841–51.

Hanna A, Metge BJ, Bailey SK, Chen D, Chandrashekar DS, Varambally S, et al. Inhibition of Hedgehog signaling reprograms the dysfunctional immune microenvironment in breast cancer. Oncoimmunology. 8; 2019.

Holla S, Stephen-Victor E, Prakhar P, Sharma M, Saha C, Udupa V, et al. Mycobacteria-responsive sonic hedgehog signaling mediates programmed death-ligand 1- and prostaglandin E2-induced regulatory T cell expansion. Sci Rep 2016;6:24193.

Hui CC, Angers S. Gli proteins in development and disease. Annual review of cell and developmental biology 2011;27:513–37.

Ikeda S, Goodman AM, Cohen PR, Jensen TJ, Ellison CK, Frampton G, et al. Metastatic basal cell carcinoma with amplification of PD-L1: exceptional response to anti-PD1 therapy. NPJ Genom Med 2016;1.

John LB, Devaud C, Duong CP, Yong CS, Beavis PA, Haynes NM, et al. Anti-PD-1 antibody therapy potently enhances the eradication of established tumors by gene-modified T cells. Clin Cancer Res 2013;19(20):5636–46.

Johnson RL, Rothman AL, Xie J, Goodrich LV, Bare JW, Bonifas JM, et al. Human homolog of patched, a candidate gene for the basal cell nevus syndrome. Science (New York, NY) 1996;272(5268):1668–71.

Kasper M, Regl G, Frischauf AM, Aberger F. GLI transcription factors: mediators of oncogenic Hedgehog signalling. Eur J Cancer 2006;42(4):437–45.

Lee JJ, Rothenberg ME, Seeley ES, Zimdahl B, Kawano S, Lu WJ, et al. Control of inflammation by stromal Hedgehog pathway activation restrains colitis. Proc Natl Acad Sci U S A 2016;113(47):E7545–E53.

Lipson EJ, Lilo MT, Ogurtsova A, Esandrio J, Xu H, Brothers P, et al. Basal cell carcinoma: PD-L1/PD-1 checkpoint expression and tumor regression after PD-1 blockade. J Immunother Cancer 2017;5:23.

Mantovani A. The yin-yang of tumor-associated neutrophils. Cancer Cell 2009;16(3):173–4.

Mittal D, Young A, Stannard K, Yong M, Teng MW, Allard B, et al. Antimetastatic effects of blocking PD-1 and the adenosine A2A receptor. Cancer research 2014;74(14):3652–8.

Mougiakakos D, Choudhury A, Lladser A, Kiessling R, Johansson CC. Regulatory T cells in cancer. Adv Cancer Res 2010;107:57–117.

Ngiow SF, Young A, Jacquelot N, Yamazaki T, Enot D, Zitvogel L, et al. A Threshold Level of Intratumor CD8+ T-cell PD1 Expression Dictates Therapeutic Response to Anti-PD1. Cancer research 2015;75(18):3800–11.

Nilsson M, Unden AB, Krause D, Malmqwist U, Raza K, Zaphiropoulos PG, et al. Induction of basal cell carcinomas and trichoepitheliomas in mice overexpressing GLI-1. Proc Natl Acad Sci U S A 2000;97(7):3438–43.

Okazaki T, Chikuma S, Iwai Y, Fagarasan S, Honjo T. A rheostat for immune responses: the unique properties of PD-1 and their advantages for clinical application. Nat Immunol 2013;14(12):1212–8.

Omland SH, Nielsen PS, Gjerdrum LM, Gniadecki R. Immunosuppressive Environment in Basal Cell Carcinoma: The Role of Regulatory T Cells. Acta dermato-venereologica 2016;96(7):917–21.

Oro AE, Higgins KM, Hu Z, Bonifas JM, Epstein EH, Jr., Scott MP. Basal cell carcinomas in mice overexpressing sonic hedgehog. Science (New York, NY) 1997;276(5313):817–21.

Papaioannou E, Yanez DC, Ross S, Lau CI, Solanki A, Chawda MM, et al. Sonic Hedgehog signaling limits atopic dermatitis via Gli2-driven immune regulation. J Clin Invest 2019;130:3153–70.

Pardoll D, Allison J. Cancer immunotherapy: breaking the barriers to harvest the crop. Nat Med 2004;10(9):887–92.

Postow MA, Chesney J, Pavlick AC, Robert C, Grossmann K, McDermott D, et al. Nivolumab and ipilimumab versus ipilimumab in untreated melanoma. The New England journal of medicine 2015;372(21):2006–17.

Robert C, Long GV, Brady B, Dutriaux C, Maio M, Mortier L, et al. Nivolumab in previously untreated melanoma without BRAF mutation. The New England journal of medicine 2015;372(4):320–30.

Ruiz i Altaba A, Mas C, Stecca B. The Gli code: an information nexus regulating cell fate, stemness and cancer. Trends Cell Biol 2007;17(9):438–47.

Samstein RM, Lee CH, Shoushtari AN, Hellmann MD, Shen R, Janjigian YY, et al. Tumor mutational load predicts survival after immunotherapy across multiple cancer types. Nat Genet 2019;51(2):202–6.

Sekulic A, Migden MR, Basset-Seguin N, Garbe C, Gesierich A, Lao CD, et al. Long-term safety and efficacy of vismodegib in patients with advanced basal cell carcinoma: final update of the pivotal ERIVANCE BCC study. BMC Cancer 2017;17(1):332.

Sekulic A, Von Hoff D. Hedgehog Pathway Inhibition. Cell 2016;164(5):831.

Sharpe HJ, Pau G, Dijkgraaf GJ, Basset-Seguin N, Modrusan Z, Januario T, et al. Genomic analysis of smoothened inhibitor resistance in basal cell carcinoma. Cancer Cell 2015;27(3):327–41.

Sternberg C, Gruber W, Eberl M, Tesanovic S, Stadler M, Elmer DP, et al. Synergistic cross-talk of hedgehog and interleukin-6 signaling drives growth of basal cell carcinoma. Int J Cancer 2018;143(11):2943–54.

Takeuchi Y, Nishikawa H. Roles of regulatory T cells in cancer immunity. International immunology 2016;28(8):401–9.

Tang JY, Mackay-Wiggan JM, Aszterbaum M, Yauch RL, Lindgren J, Chang K, et al. Inhibiting the hedgehog pathway in patients with the basal-cell nevus syndrome. The New England journal of medicine 2012;366(23):2180–8.

Teglund S, Toftgard R. Hedgehog beyond medulloblastoma and basal cell carcinoma. Biochimica et biophysica acta 2010;1805(2):181–208.

Wei SC, Duffy CR, Allison JP. Fundamental Mechanisms of Immune Checkpoint Blockade Therapy. Cancer Discov 2018;8(9):1069–86.

Whitson RJ, Lee A, Urman NM, Mirza A, Yao CY, Brown AS, et al. Noncanonical hedgehog pathway activation through SRF-MKL1 promotes drug resistance in basal cell carcinomas. Nat Med 2018;24(3):271–81.

Xie J, Murone M, Luoh SM, Ryan A, Gu Q, Zhang C, et al. Activating Smoothened mutations in sporadic basal-cell carcinoma. Nature 1998;391(6662):90–2.

Zhou X, Liu Z, Jang F, Xiang C, Li Y, He Y. Autocrine Sonic hedgehog attenuates inflammation in cerulein-induced acute pancreatitis in mice via upregulation of IL-10. PLoS One 2012;7(8):e44121.

