## Supplementary Materials and Methods for "Epidermal activation of Hedgehog signaling establishes an immunosuppressive microenvironment in basal cell carcinoma by modulating skin immunity"

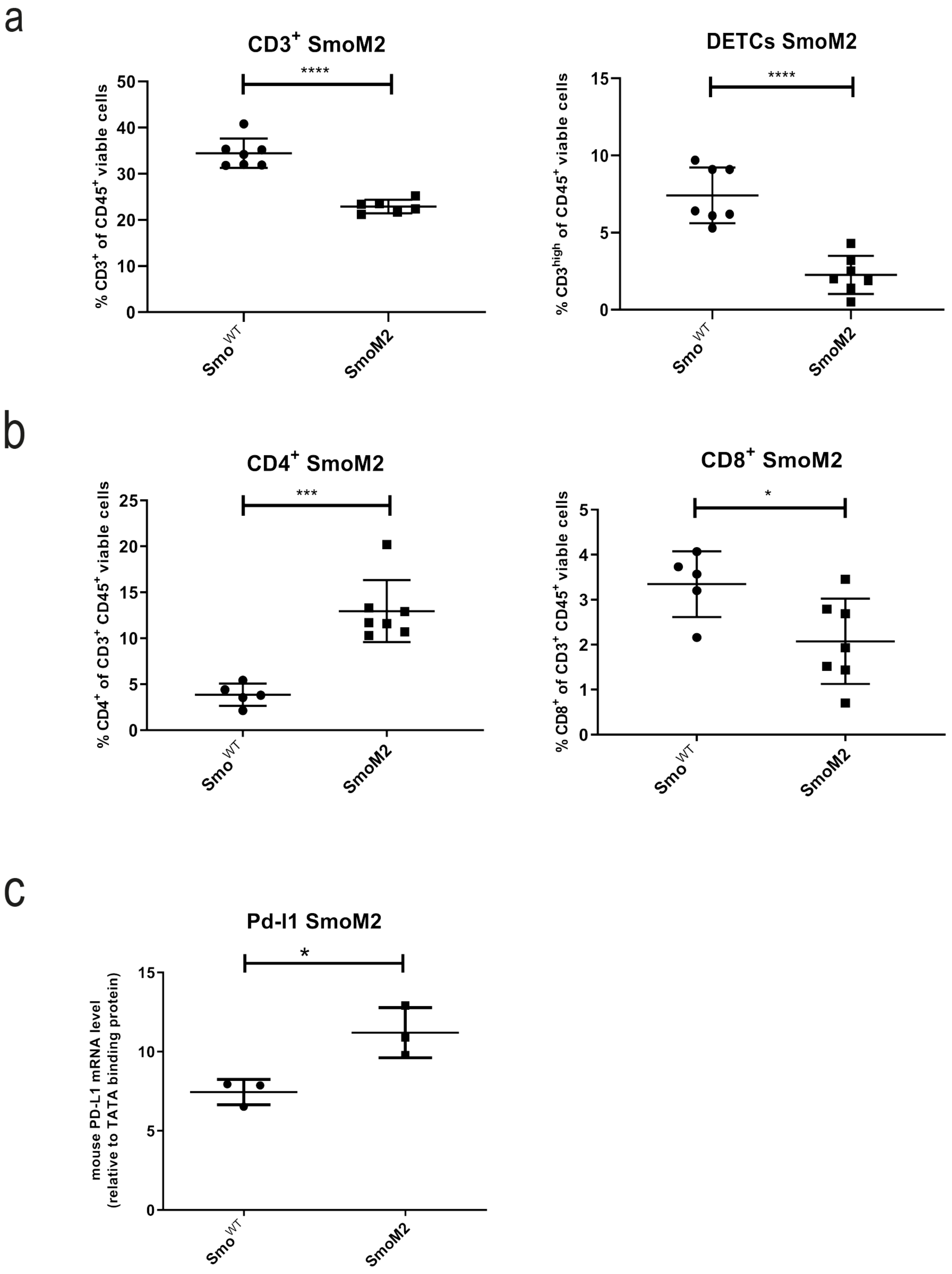


Grund-Groeschke et al. Figure S1

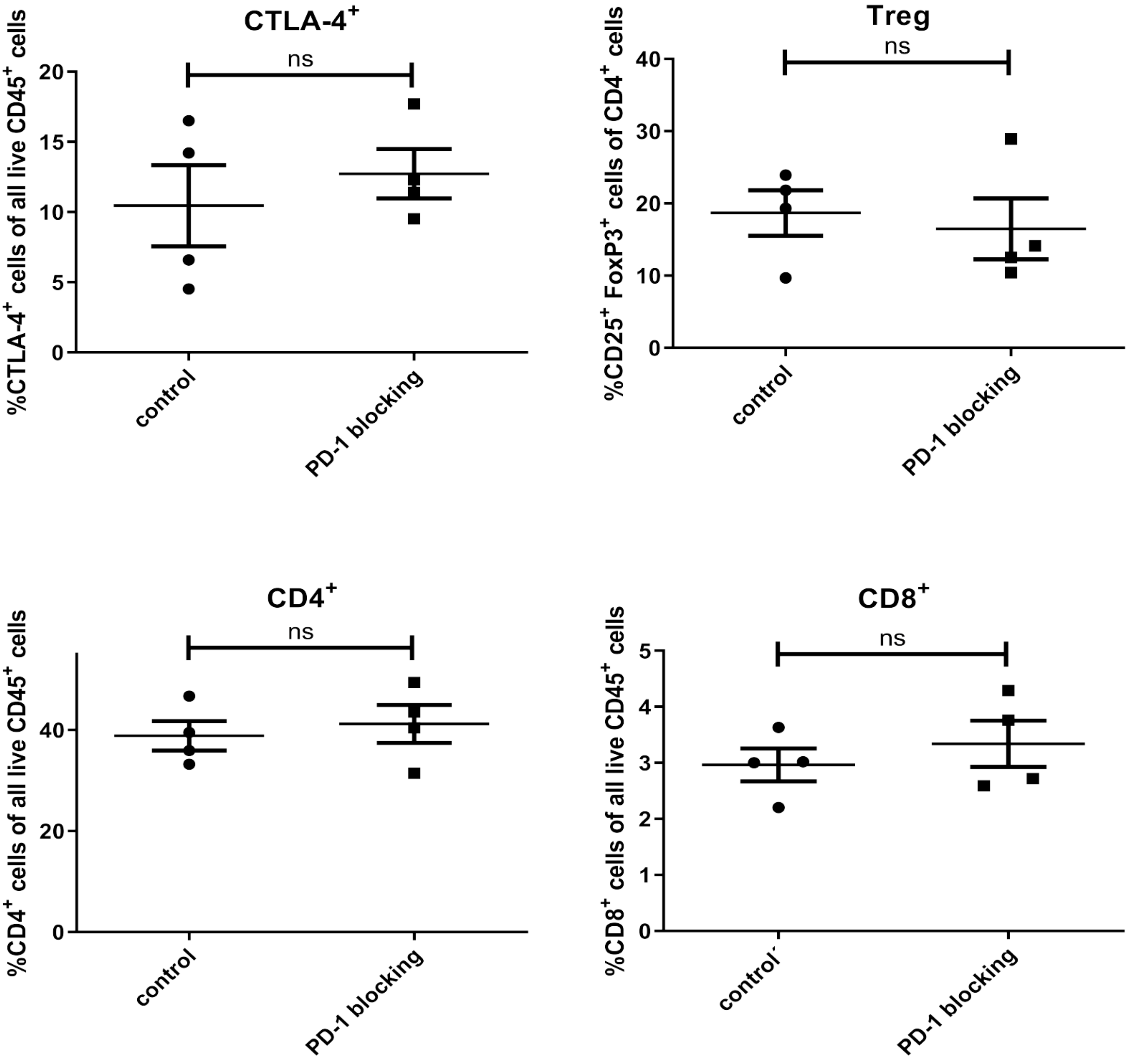


Grund-Groeschke et al. Figure S2

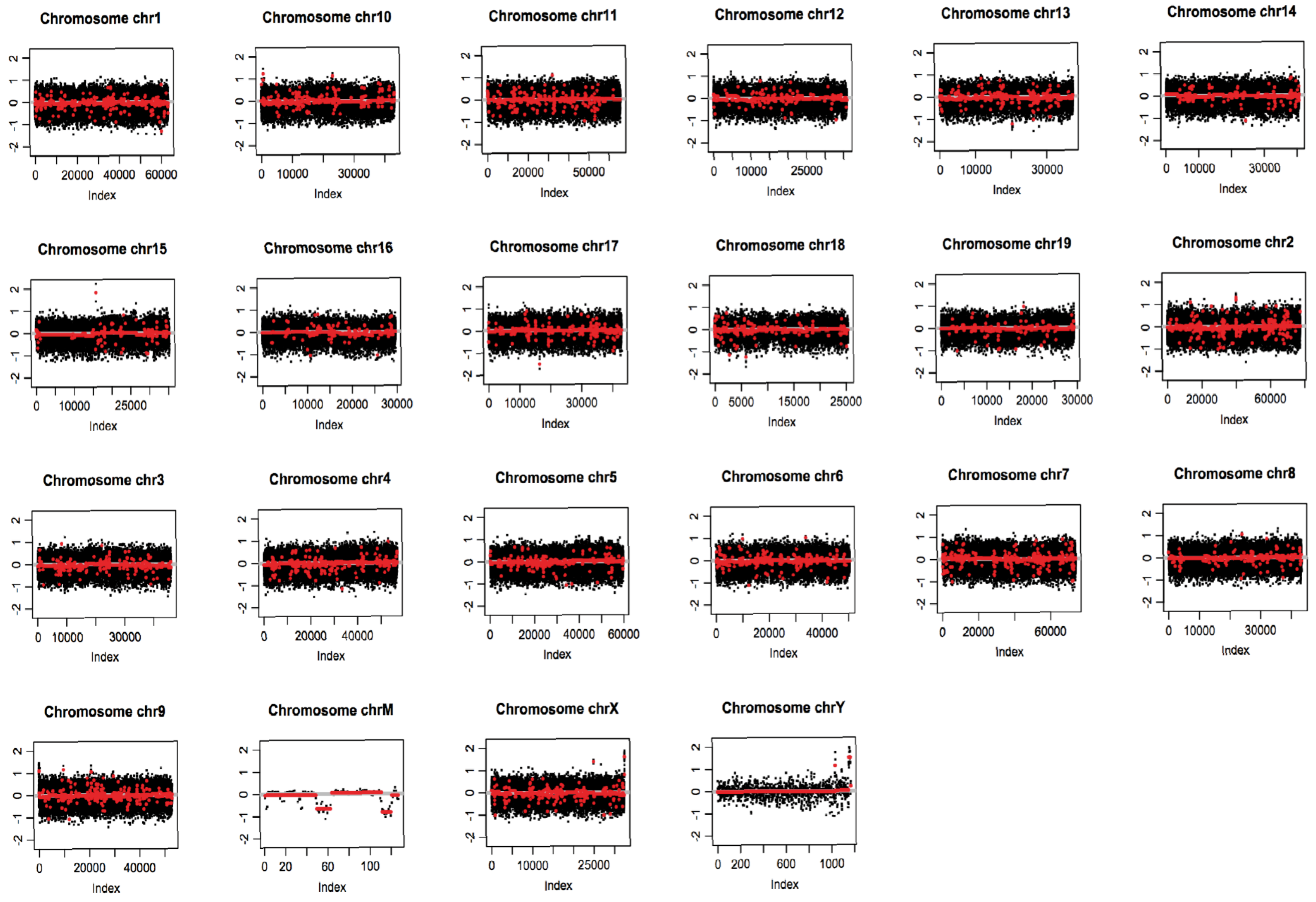


Grund-Groeschke et al. Figure S3

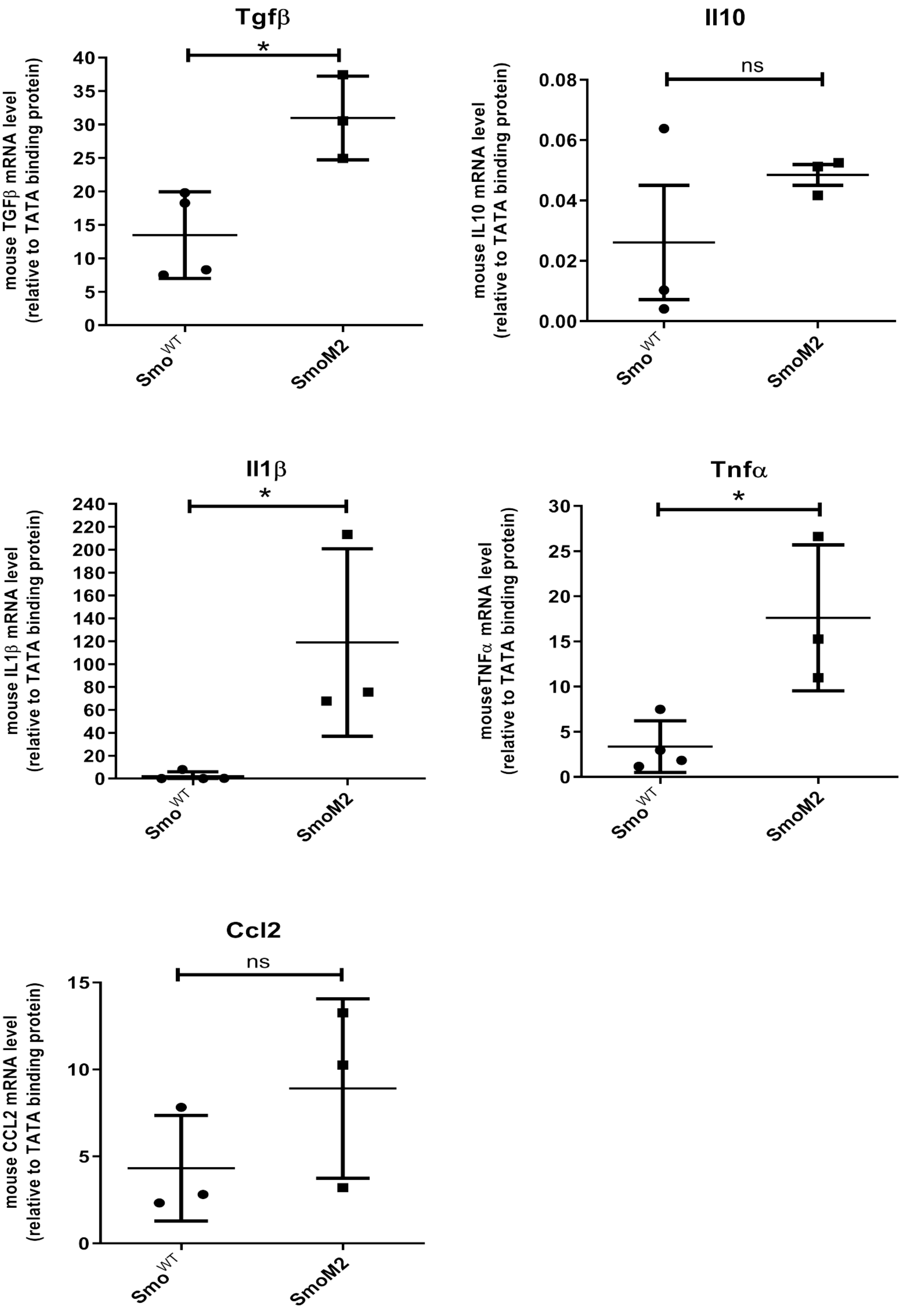


Grund-Groeschke et al. Figure S4

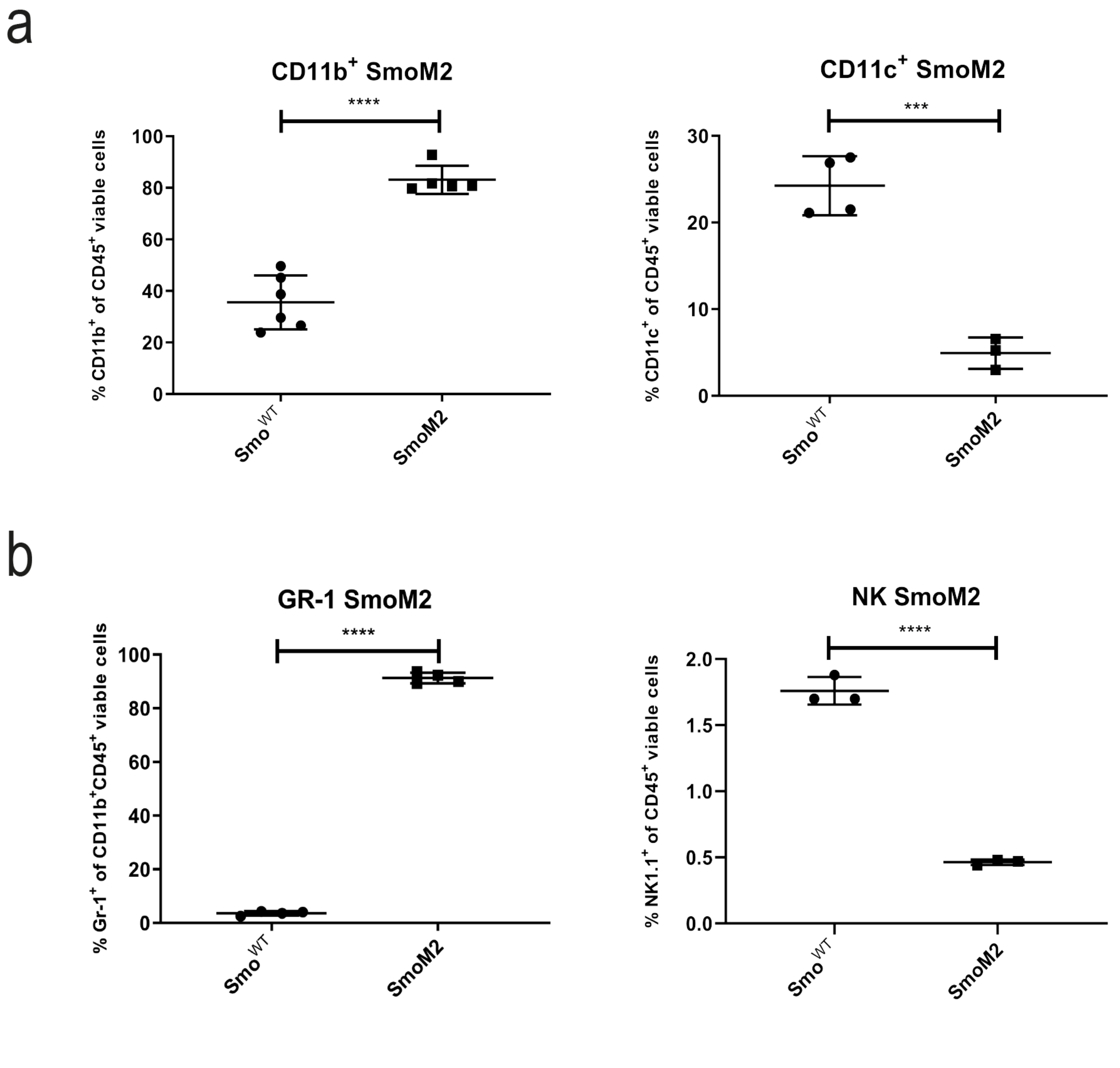


Grund-Gröschke et al. Figure S5

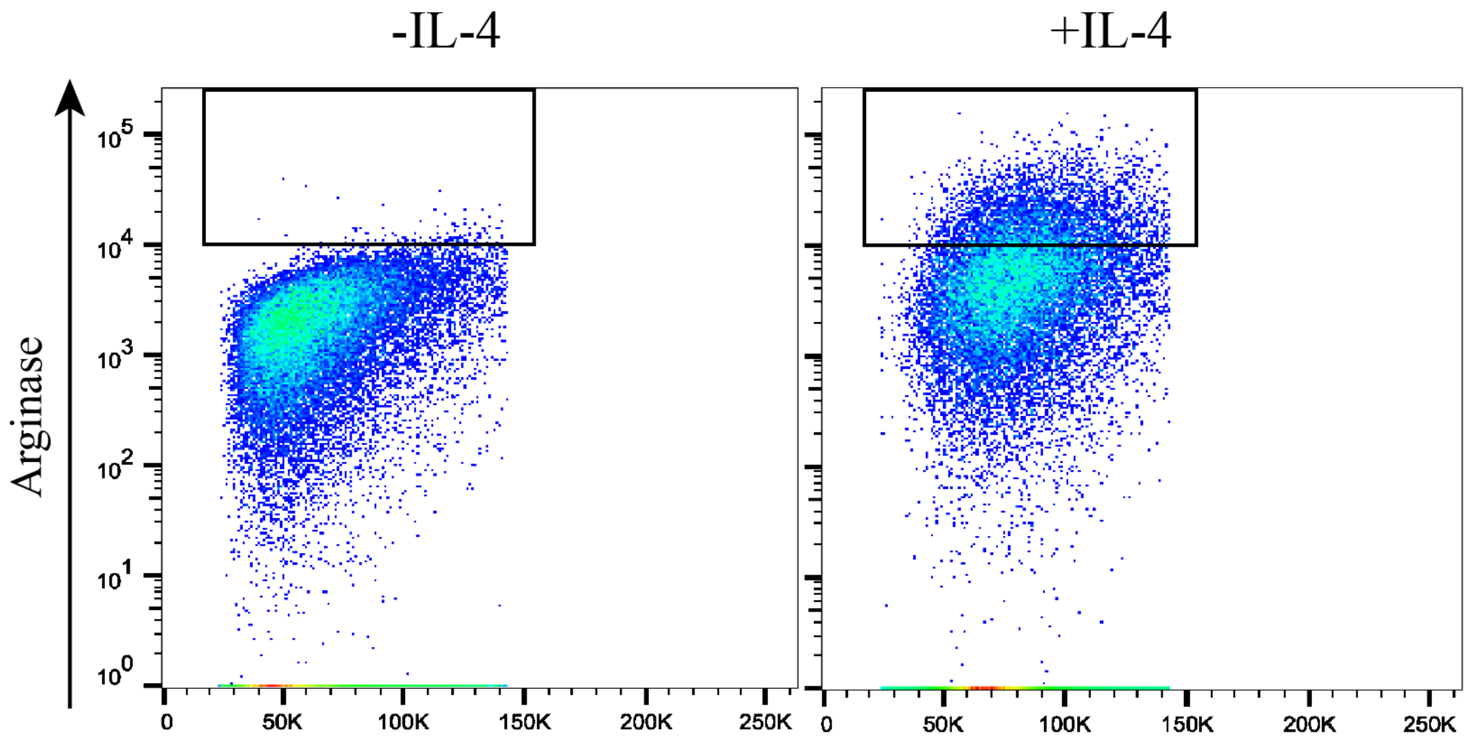


Grund-Gröschke et al. Figure S6
