## Supplementary Figures S1-S6 for "Epidermal activation of Hedgehog signaling establishes an immunosuppressive microenvironment in basal cell carcinoma by modulating skin immunity"

### Supplementary Material:

#### Supplementary Tables

| Suppl. Table S1: Mutational landscape of tumors from *Ptch*$^{\Delta ep}$ mice* | | | | | |
|---|---|---|---|---|---|
| **chromosome** | **Position** | **mutation type** | **Mutation** | **consequence** | **Gene** |
| chr6 | 91504069 | base exchange | C -> G | D274H | Xpc |
| chr18 | 24601227 - 24601244 | deletion | TGCTCGTGGTC CGAGTGA | frame shift | Slc39a6 |

* Table shows mutations found by whole exome sequencing in one out of three *Ptch*$^{\Delta ep}$ mice. Chr= chromosome.

| Suppl. Table S2: Antibodies used for flow cytometry | | | | |
|---|---|---|---|---|
| **Antibody** | **Clone** | **Company** | **Dilution** | **Identifer** |
| CD45 | 30-F11 | Thermo Scientific | 1:1500 | 47-0451 |
| CD11b | M1/70 | Thermo Scientific | 1:500 | 69-0112 |
| Ly-6G | 1A8 | Thermo Scientific | 1:500 | 17-9668 |
| CD11c | N418 | Thermo Scientific | 1:200 | 25-0114 |
| NK-1.1 | PK136 | Thermo Scientific | 1:200 | 12-5941 |
| CD3 | 145-2C11 | Thermo Scientific | 1:300 | 45-0031 |
| CD4 | GK1.5 | Thermo Scientific | 1:200 | 11-0041 |
| CD4 | GK1.5 | Thermo Scientific | 1:2000 | 12-0041 |
| CD8a | 53-6.7 | Thermo Scientific | 1:400 | 25-0081 |
| CD8a | 53-6.7 | Tonbo Biosciences | 1:400 | 20-0081 |
| γδTCR | eBioGL3 | Thermo Scientific | 1:600 | 12-5711 |
| FoxP3 | FJK-16s | Thermo Scientific | 1:300 | 48-5773 |
| CD274/PD-L1 | MIH5 | Thermo Scientific | 1:500 | 12-5982 |
| CD49f | eBioH3 | Thermo Scientific | 1:500 | 25-0495 |
| Ly-6A/E | D7 | Thermo Scientific | 1:1000 | 62-5981 |
| CD279/PD-1 | J43 | Thermo Scientific | 1:200 | 11-9985 |
| CD25 | PC61.5 | Tonbo Biosciences | 1:500 | 12-0251 |
| Arginase | polyclonal | R&D Systems | 1:20 | IC5868P |

| | | | | |
|---|---|---|---|---|
| CD45 | 30-F11 | Biolegend | 1:100 | 103138 |
| CD4 | GK1.5 | Biolegend | 1:100 | 100408 |
| CD8 | 53-6.7 | Biolegend | 1:100 | 100752 |
| CD3 | 17A2 | Biolegend | 1:100 | 100234 |
| CD11c | N418 | Biolegend | 1:100 | 117318 |
| Gr1 | RB6-8C5 | BD | 1:2500 | 562709 |
| CD11b | M1/70 | Biolegend | 1:200 | 101228 |
| NK1.1 | PK136 | Biolegend | 1:100 | 108728 |
| CD45 | 30-F11 | Biolegend | 1:100 | 103138 |

**Suppl. Table S3: Primer sequences used for qPCR of murine samples**

| Gene name | Forward primer (5’-3’) | Reverse primer (5’-3’) |
|---|---|---|
| Rplp0 | ATAACCCTGAAGTGCTCGACAT | CCATTGATGATGGAGTGTGG |
| Gli1 | CACCGTGGGAGTAAACAGGCCTTCC | CCAGAGCGTTACACACCTGCCCTTC |
| Gli1 | CACATCAACAGTGAGCATATCCA | GTGAATAGGACTTCCGACAGC |
| Il17 | TCTCTGATGCTGTTGCTGCTGCTGA | GTCCAGCTTTCCCTCCGCATTGAC |
| Il10 | CTCCTAGAGCTGCGGACTGCCTTCA | CTTCACCTGCTCCACTGCCTTGCTC |
| Tgf-b | ATGAACCGGCCCTTCCTGCTCCT | CAGAAGTTGGCATGGTAGCCCTTGG |
| Nos2 | CCTGCTTTGTGCGAAGTGTCAGTGG | TCTCTTGCGGACCATCTCCTGCATT |
| Il1b | TACAAGGAGAACCAAGCAACGACAAAATACC | AGGGTGGGTGTGCCGTCTTTCAT |
| Ifng | TGAAAGACAATCAGGCCATCAGCA | CAGCAGCGACTCCTTTTCCGCTTC |
| Ccl2 | CCAGCTCTCTCTTCCTCCACCACCA | ACCCATTCCTTCTTGGGGTCAGCAC |
| Ccl3 | GCAGCAGCGAGTACCAGTCCCTTTTC | TCTTCCGGCTGTAGGAGAAGCAGCA |
| Pd1 | GCAATCAGGGTGGCTTCTA | GTTCCAGTTCAGCATAAGATCCTC |
| Pdl1 | CCGGACAGAGGGGATGCTTCTCA | TGTGGAGGATGTGTTGCAGGCAGTT |
| Pdl2 | GCAAAGTGAAAGAGCCACCCTGCTG | GCTGCACCTCCCCTGTACCTGGA |
| Tim3 | TACAGTTCCCTGGTCTTATGAATGA | GTCTGTGTCTCTGAACCATTTCTCT |
| Tigit | ATGGCTGCTGTGCTGGGACTCATTT | TTCCATTCCTGTGGCTCCGCTTCT |
| Lag3 | ACACCTGTAGCATCCATCTGC | CCACACAAATCTTTCCTTTCCAG |
| Cd226 | TCCTGCTTGTTTCATGCTTTCCCAAAT | TCTACCTGATGGGGCTGGACTTTTTCC |
| Cd96 | ATACCATCATCAGTACAACCACAGA | ACCGATACCATTTTCTTACTCCAAG |

| Suppl. Table S4: Antibodies used for immunofluorescence | | | |
|---|---|---|---|
| **Antibodies** | **Source** | **Identifier** | **Dilution** |
| Rabbit anti-PD-1 | Cell Signaling | 84651 | 1:100 |
| Rabbit anti-FoxP3 | Cell Signaling | 12653 | 1:100 |
| Rat anti-Ly6G | Biolegend | 127601 | 1:1000 |
| Rabbit anti-CD8 | Cell Signaling | 98941 | 1:200 |
| Goat anti-Rabbit IgG (H+L), Alexa Fluor 488 | Thermo Fisher | A-11008 | 1:1000 |
| Goat anti-Rat IgG (H+L), Alexa Fluor 555 | Thermo Fisher | A-21434 | 1:1000 |

| Suppl. Table S5: qPCR primer list for *SmoM2* mice: | |
|---|---|
| **Gene id** | **Assay number *** |
| Il10 | Mm00439614_m1 |
| Tnf | Mm00443258_m1 |
| Ccl2 | Mm00441242_m1 |
| Tgfb | Mm01178820_m1 |
| Il1b | Mm00434228_m1 |
| Pd-l1 | Mm03048248_m1 |

* All primers for analysis of *SmoM2* mice were obtained from Thermo Scientific/Applied Biosystems.

### Supplementary Methods

#### Mice

*R26SmoM2:YFP* (Mao et al, 2006) and *K5cre$^{ER}$* (Ramirez et al, 2004) mice were breed to obtain *K5cre$^{ER}$*;R26*SmoM2:YFP (*SmoM2) mice. Activation of *SmoM2* expression was accomplished by i.p. injection of 0.5 mg tamoxifen citrate per day starting at postnatal day 14 for 5 consecutive days; Cre negative littermates received the same treatment. Mice started to develop macroscopically visible tumors 5 weeks after tamoxifen administration.

#### Flow cytometry

For flow cytometry analysis of *SmoM2* mice the ear skin was cut into small pieces and digested with 0.15 mg/ml Liberase™ and 0.12 mg/ml DNAse I (both Roche Diagnostics, Indianapolis, IN) for 45 minutes at 37°C and pressed through 100 µm cell strainers (BD Biosciences, Franklin Lakes, NJ). Nonspecific FcR-mediated antibody staining was blocked with anti-CD16/32 (2.4G2, in-house from hybridoma supernatant or from BD Biosciences, Franklin Lakes, NJ) for 15 minutes at 4°C. Cells were stained with directly conjugated antibodies for 15 min at 4°C in the dark. Dead cells were excluded with 7-AAD (BD Biosciences, Franklin Lakes, NJ) or fixable viability dye eFluor® 780 (Thermo Fisher Scientific, Waltham, MA). To separate immune cells from other skin cells CD45 as pan-leukocyte marker was included in all skin panels. Cells were fixed by using the Foxp3/transcription factor staining buffer set (Thermo Fisher Scientific, Waltham, MA) or the Cytofix/Cytoperm™ kit (BD Biosciences, Franklin Lakes, NJ) followed by intracellular staining. All experiments were performed on the BD FACS Canto II (BD Biosciences, Franklin Lakes, NJ). For data analysis the FlowJo® software (Tree Star) was used. Antibodies used for flow cytometry are listed in Supplement Table S2.

#### Quantitative PCR

Total RNA from *SmoM2* mouse skin was isolated with TRIZOL (Thermo Fisher Scientific, Waltham, MA). Genomic DNA was removed using the RapidOut DNA removal kit (Thermo

Fisher Scientific, Waltham, MA). Random primed cDNA was prepared with the Superscript II RNase H-reverse transcriptase (Thermo Fisher Scientific, Waltham, MA). Cytokine mRNA expression were examined with quantitative polymerase chain reaction (PCR) analysis by using the Brilliant III Ultra-fast qPCR Master Mix (Agilent Technologies, Santa Clara, CA). Information about the primers used is listed in Suppl. Table S5.

**Mouse whole exome sequencing**

Mouse genomic DNA was isolated from ear tissue using the DNeasy Blood and Tissue kit (Qiagen, Hilden, Germany). Isolated genomic DNA was sheared with the Covaris M220 (Covaris, Woburn, MA). DNA was subjected to whole exome library generation (SureSelect$^{XT}$ Mouse All Exon Kit; 49.6 megabases (Agilent, Santa Clara, CA)), which were sequenced 100 bp paired-end on the Illumina platform NextSeq 550 using the NextSeq 500/550 v2.5 kit (Illumina, San Diego, CA). Sequencing reads were mapped to mouse reference genome (UCSC mm10) using Burrows-Wheeler Aligner with default settings (BWA-MEM v0.7.12-R1039) (Li and Durbin, 2010). Duplicate removal was performed with default parameters using PicardTools (v2.10.3, http://broadinstitute.github.io/picard/). Alignments were preprocessed including local realignment around indels and base quality recalibration was performed using Genome Analysis Tool Kit (GATKv3.7) (all with default parameters) (McKenna et al., 2010). For somatic mutation calling (SNVs, Indels) each tumor sample (*K14CreER$^{T}$;Ptch$^{f/f}$*) was compared with three *Ptch$^{f/f}$* mice. Subsequently, all unique SNVs and indels were called. Mpileup file generation was done by samtools (v1.5) (parameter –B –q 1) (Li et al., 2009), VarScan2 (v2.4.3) was used for somatic variant calling (–min-coverage-normal 5 –min-coverage-tumor 5 –min-var-freq 0.05 -somatic-p-value 0.05 –strand-filter 1) and filtering of high confidence calls was performed according to Basic Protocol 2 published by Koboldt et al. (Koboldt et al., 2013). Variants were annotated using ANNOVAR (version 2017Jul16) (Wang et al., 2010). All programs were executed following the authors' recommendations. Nonsynonymous exonic SNVs and Indels were considered. All remaining mutations were individually checked for accuracy using Integrative Genomics Viewer (IGV) version 2.4.2. (Robinson et al., 2011, Thorvaldsdottir et al., 2013). For Copy Number Variations (CNVs) detection, depth of coverage was calculated for each exome target region using Varscan copynumber and Varscan copycaller. Coverage data was then analyzed using the R (version 3.6.1) package DNACopy (version 1.58.0) using the circular binary segmentation algorithm (CBS) to segment DNA copy number data and identify genomic regions with abnormal copy number (Seshan and Olshen, 2019).

**Alternative activation of mouse bone marrow-derived macrophages**

Mouse bone marrow cells were isolated from the femoral bones and after red blood cell lysis grown in L929 fibroblast-conditioned medium (L929-sup, made in house) for ten days at 7%$CO_2$. Afterwards, bone marrow macrophages were alternatively activated with 50 ng/ml recombinant mouse IL4 (Immunotools, Friesoythe, Germany) for 24h.

### Supplementary Figure legend:

**Figure S1: Altered immune phenotype of *SmoM2* mice.** (a-d) Flow cytometry analysis of (a) $CD3^{+}$ and $CD3^{high}$ T cells, (b) CD4 and CD8 T cells in $Smo^{WT}$ and *SmoM2* mice. (c) mRNA expression level of *Pd-l1* in $Smo^{WT}$ and *SmoM2* mice.

**Figure S2: Anti-Pd-1 blocking does not reduce skin cancer phenotype of $Patched^{\Delta ep}$ mice.** Flow cytometry analysis of the skin after Pd-1 blocking.

**Figure S3: Mouse tumors exhibit no large structural genetic variations.** Representative copy number variation (CNV) plot from mouse #2. CNV analysis reveals no deletions or amplifications.

**Figure S4: Altered cytokine and chemokine profile of *SmoM2*.** Cytokine and chemokine profiles were determined via mRNA expression level of $Smo^{WT}$ and *SmoM2* mice.

**Figure S5: Altered innate immunity in *SmoM2* mice.** (a-d) Flow cytometry analysis of (a) CD11b and CD11c innate immune cells and (b) $GR\text{-}1^{+}$ and $NK1.1^{+}$ cells in $Smo^{WT}$ and *SmoM2* mice.

**Figure S6: Validation of the arginase antibody.** Representative flow cytometry plots of bone marrow-derived macrophages unstimulated or stimulated with Il4, a known inducer of arginase expression.

**References:**


Koboldt DC, Larson DE, Wilson RK. Using VarScan 2 for Germline Variant Calling and Somatic Mutation Detection. Curr Protoc Bioinformatics 2013;44:15.4.1-7.

Li H, Durbin R. Fast and accurate long-read alignment with Burrows-Wheeler transform. Bioinformatics 2010;26(5):589-95.

Li H, Handsaker B, Wysoker A, Fennell T, Ruan J, Homer N, et al. The Sequence Alignment/Map format and SAMtools. Bioinformatics 2009;25(16):2078-9.

McKenna A, Hanna M, Banks E, Sivachenko A, Cibulskis K, Kernytsky A, et al. The Genome Analysis Toolkit: A MapReduce framework for analyzing next-generation DNA sequencing data. Genome Res 2010;20(9):1297-303.

Robinson JT, Thorvaldsdottir H, Winckler W, Guttman M, Lander ES, Getz G, et al. Integrative genomics viewer. NatBiotechnol 2011;29(1):24-6.

Seshan V, Olshen A. DNAcopy: DNA copy number data analysis. R package 2019;version 1.58.0.

Thorvaldsdottir H, Robinson JT, Mesirov JP. Integrative Genomics Viewer (IGV): high-performance genomics data visualization and exploration. Brief Bioinform 2013;14(2):178-92.

Wang K, Li M, Hakonarson H. ANNOVAR: functional annotation of genetic variants from high-throughput sequencing data. Nucleic Acids Res 2010;38(16):e164.
